# Differences in perceived duration between plausible biological and non-biological stimuli

**DOI:** 10.1101/664193

**Authors:** Giuliana Martinatti Giorjiani, Claudinei Eduardo Biazoli, Marcelo S. Caetano

## Abstract

Visual motion stimuli can sometimes distort our perception of time. This effect is dependent on the apparent speed of the moving stimulus, where faster stimuli are usually perceived lasting longer than slower stimuli. Although it has been shown that neural and cognitive processing of biological motion stimuli differ from non-biological motion stimuli, no study has yet investigated whether perceived durations of biological stimuli differ from non-biological stimuli across different speeds. Here, a prospective temporal reproduction task was used to assess that question. Biological motion stimuli consisted in a human silhouette running in place. Non-biological motion stimuli consisted in a rectangle moving in a pendular way. Amount and plausibility of movement for each stimulus and frame-rate (speed) were evaluated by an independent group of participants. Although amount of movement was positively correlated to frame rate, movie clips involving biological motion stimuli were judged to last longer than non-biological motion stimuli only at frame rates in which movement was rated as plausible. These results suggest that plausible representations of biomechanical movement induce additional temporal distortions to those modulated by increases in stimulus speed. Moreover, most studies that have reported neural and cognitive differences in the processing of biological and non-biological motion stimuli acquired neurophysiological data using fMRI. The present study aimed additionally to report differences in the processing of biological and non-biological motion stimuli across different speeds using functional near infrared spectroscopy (fNIRS), a less costly and portable form of neurophysiological data acquisition.

## Introduction

The perception of time is ubiquitous in nature and an important part of most human behaviors. From planning daily routines to executing specific motor responses, many activities can be summarized as a series of successive events in time. Despite our ability to perceive the passage of time and interact with the environment accordingly, different factors can warp our sense of time, such as attention, arousal, temporal frequency and motion perception [1-4]. Motion stimuli in particular are especially useful in the study of temporal distortions, as the perception of natural motion relies on a smooth sequence of events occurring in a time array [5]. Additionally, when represented virtually, natural motion depends on discrete events being interspersed by an optimal interval (i.e., optimal frame rate [6]).

Previous studies have all shown that distortions in temporal estimation of moving stimuli are speed-dependent [7-10]. Their results suggest that the higher the speed of a moving stimulus, the longer the interval is perceived. In general, durations of low speed stimuli are underestimated compared to high speed stimuli. Different accounts of such results include speeding up of a pacemaker in internal clock models [7, 11, 12], and an effect of the number of spatial changes (higher for high-speed stimuli compared to low-speed stimuli) leading to temporal overestimation [8, 13]. Alternatively, instead of a single cognitive timing mechanism, multiple mechanisms might be in place [14]. Building upon results regarding visual processing in the sub-second scale, Bruno and Cicchini proposed that several parallel clocks might be instantiated at multiple points of the visual pathway [14]. Considering that the spatial features and speed of moving stimuli can differently recruit high-order visual areas (e.g., in the illusion of movement depending on frame rates), the existence of multiple timing mechanisms might lead to differences in temporal perception of such stimuli.

To account for speed-dependent effects on time perception, Fraisse has proposed that an increase in the number of observed distinguishable changes would lead to dilation in temporal perception [5]. Specifically, she argues that the greater the frame rate of static visual stimuli, the more overestimated will be the time interval. However, above a certain threshold, higher frame rates will induce the illusion of fluid rather than segmented movement, possibly canceling any temporal dilation.

Accordingly, Nyman and colleagues have found no differences in time reproduction for frame rates of 25 and 50 frames per second when tested with different movement-speeds in video sequences [15]. However, to best of our knowledge, no systematic evaluations of temporal distortions by frame rates above and below induction of motion illusion have been conducted. Potential effects of plausible biological movement stimuli on time reproduction for different frame rates and related perceived speed have also not been comprehensively assessed.

Most of previous studies addressing effects of movement on time perception have used geometric shapes moving across a computer screen as their primary stimuli. However, static and dynamic representations of the human body can also generate temporal distortions [16, 17]. Apparent biological motion, particularly regarding the shape of the human body, is thought to induce specific distortions in time perception [18-23]. Nather and Bueno suggested that the perceived duration of symmetrical representations of the human body (sculptures) tended to be temporally underestimated compared to asymmetrical body sculptures [16]. According to the authors, asymmetric figures implicitly suggest a larger amount of movement, as they require larger and longer actions to reestablish the body balance compared to symmetrical figures. Orgs and colleagues have shown that a sequence of frames representing a dynamic human movement can also distort time perception [17]. In their study, however, the higher the perceived speed of the human body, the shorter the perceived duration of the movement.

Besides exhibiting different behavioral properties, differences between biological and non-biological motion stimuli are also found in terms of cognitive and neural processing. Several studies suggest that the perception of visual biological motion recruits different high-level visual processing when compared to non-biological motion stimuli [22-26]. Remarkably, neurons in ventral premotor and inferior parietal cortices were shown to be activated by the observation of biological movement [27]. In accordance with findings with non-human animals, fMRI studies in humans have also suggested that different cortical areas are involved in the processing of biological vs. non-biological motion, with superior temporal sulcus (STS) preferably encoding biological motion while middle temporal gyrus (MTG) correlated with observing non-biological motion [28-34]. These results were corroborated by Ishizy and colleagues, who have used functional near infrared spectroscopy (fNIRS) to show higher hemodynamic activity in the extrastriate body area (EBA), which sits close to STS, when participants were presented with human-like figures [35].

Given the specificity in the neural processing of visual stimuli depicting human motion – and the hypothesis of multiple channels for timing perception – we tested whether a simplified representation of a moving human body (i.e., silhouette) would induce a distortion in temporal perception when compared to a non-human moving figure (i.e., geometric form). Critically, given the well-recognized speed-dependent effects of frame rate on time perception, a range of frame rates was tested, which were also evaluated in terms of amount and plausibility of movement they produced.

## General materials and methods

### Stimuli

In all experiments herein described, participants watched a series of movie clips composed of biological or non-biological stimuli. Biological motion stimuli consisted of a stroboscopic movie clip composed of 12 frames. Each frame showed a two dimensional, 8-cm tall representation of a running human silhouette at the center of the computer screen. Non-biological motion stimuli also consisted of a stroboscopic movie clip composed of 12 frames each with an 8×2 cm rectangular bar, which moved in pendular way and was presented at the center of the screen. Images of each frame from biological and non-biological motion stimuli were approximately equated by size, color and between-frames displacement. Biological and non-biological movie clips always lasted 15 seconds but were presented in five different frame rates: 0.2, 0.4, 6.4, 12.8, and 25.6 frames per second (fps).

To keep the duration of each movie clip constant across the different frame rates, the number of frames composing each movie clip was manipulated. For example, the number of frames in the 0.2 fps movie clip was 3 (0.2 × 15), while the number of frames in the 25.6 fps movie clip was 384 (25.6 × 15). If the number of frames in the movie clip was lower than 12 (the original set of frames), intermediate frames were used (e.g., for the 0.2 fps, frames 1, 6, and 12 were used). If the number of frames in the movie clip was higher than 12, the whole set of frames was repeatedly presented until the desired duration was achieved (e.g., for the 25.6 fps, the set of 12 frames was presented 32 times, totaling 384 frames).

Stimuli were displayed on a 24 inches DELL LED screen (64 Hz refresh rate) using a DELL Optiplex 9010 desktop computer. All experimental events (stimuli presentation, recording of responses etc.) were controlled by a MATLAB Psychtoolbox script. All experimental protocols in the three experiments herein reported were approved by the Institutional Review Board of Federal University of ABC.

### Temporal reproduction task

In Experiments II and III, participants performed a temporal reproduction task similar to that used by Brown, Kanai, Nather and Bueno [7,8,16]. They were seated 60 cm from the computer monitor and informed about the general task procedure. They were told to pay close attention to the movie clips and then reproduce their perceived durations.

Each trial began with the presentation of one of the 10 possible movie clips for 15 seconds. Next, an instruction screen displayed the following message: “Press the A key to start the temporal reproduction and the L key to end it”. When the participant pressed the A key, a screen with a centered cross target was presented until the participant pressed the L key to end the interval reproduction. The next screen showed the following message: “Press SPACEBAR to continue”. A press to the space bar started the next trial (Figure 2). Each participant was trained with 10 blocks of trials. Each block was composed of a random permutation of the 10 trial types. Therefore, by the end of the experiment, each participant reproduced each of the 10 trials types 10 times.

Descriptive and inferential analyses of performance in this task were performed on a 20% trimmed mean of the reproduced intervals for each of the 10 trial types (i.e., the shortest and highest intervals reproduced were excluded). Individual data were then submitted to a repeated measures general linear model analysis (GLM) and Wilcoxon tests for paired samples.

## Experiment I – Motion stimuli evaluation

The goal of Experiment I was to assess the perceived amount of movement and plausibility for each of the 10 movie clips. Specifically, this experiment sought to confirm that higher frame rates correlated to increasing perceived speeds, and to determine what frame rates were judged as representing natural smooth movements.

### Participants

Seventeen healthy undergraduate students participated in this experiment (8 women, mean age: 23.0 ±3.7 years).

### Procedure

Participants were seated 60 cm from the computer monitor and informed about the general task procedure. They were told to pay close attention to the movie clips and report (on a paper form) amount and plausibility of movement after each trial. They had to judge both “how much movement” they observed and whether the moving object seemed natural (see Supporting Information). Each trial began with the presentation of one of the 10 possible movie clips for 15 seconds (biological and non-biological stimuli, each presented at the five possible frame rates). Next, they reported on a 10-point scale how much movement was involved in the sequence just observed, 0 (little movement) to 10 (high movement), and whether the movement seemed natural or non-natural (binary response). A press on the space bar started the next trial. The experimental session consisted of three blocks of trials. Each block was composed of a random permutation of the 10 trial types. Therefore, by the end of the experiment, each participant evaluated each of the 10 trial types three times. Before the first block of trials, participants practiced with three sample trials each with a five-second movie clip of twelve frames displaying different geometric forms.

### Analysis

Average scores for amount of movement and plausibility of movement were calculated considering the evaluations for the second and third blocks of trials (i.e., ratings for the first time they observed each movie clip were not considered). Binary choices for plausibility were computed as 0 (plausible) and −1 (not plausible). A repeated measures general linear model analysis (GLM) was performed on data for plausibility and amount of movement. To further examine the effects of frame rate on perceived amount of movement, Spearman correlations and Wilcoxon tests for paired samples were used.

### Results

Judgments of amount of movement varied as a function of frame rate, but not as a function of stimulus type (Figure 1A). For the biological stimuli, mean (± SEM) estimates for the amount of movement were 1.32 (± 0.22), 2.20 (± 0.40), 4.32 (± 0.35), 5.94 (± 0.33), and 8.53 (± 0.22) for frame rates 0.2, 0.4, 6.4, 12.8, and 25.6, respectively. For the non-biological stimuli, mean (± SEM) estimates for the amount of movement were 1.67 (±0.45), 2.76 (± 0.29), 4.64 (± 0.24), 5.97 (± 0.35), and 8.26 (±0.28) for frame rates 0.2, 0.4, 6.4, 12.8, and 25.6, respectively. A GLM analysis with two factors, type of stimulus (biological and non-biological) and frame rate (0.2; 0.4; 6.4; 12.8, and 25.6 fps) showed an effect of frame rate (F(4,64) = 167.64, p < 0.001), no effect of type of stimulus (F(1,16) = 0.642, p = 0.435) and no type of stimulus × frame rate interaction (F(4,64) = 0.86, p = 0.494). Individual Spearman correlations for biological (*ŕ* = 0.93, Z = 3.63, p < 0.001) and non-biological stimuli (*ŕ* = 0.89, Z = 3.62, p < 0.001) suggested that judged amount of movement increased linearly with frame rate.

**Figure 1.**
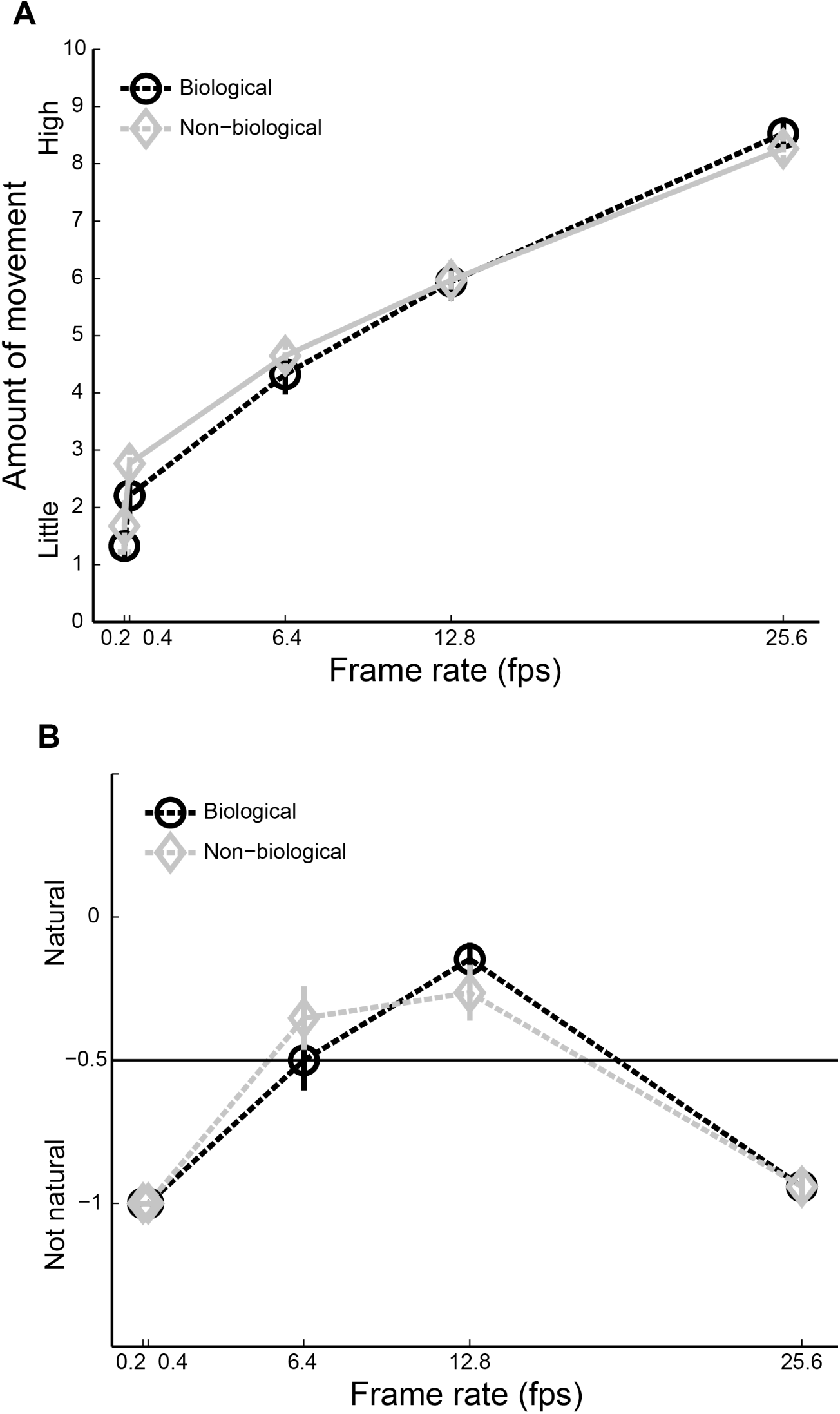
Amount of movement and plausibility assessment. (A) Mean amount of movement reported as a function of frame rate for biological (black) and non-biological (gray) movie clips. (B) Mean judgement of plausibility as a function of frame rate for biological (black) and non-biological (gray) movie clips. The horizontal line at −0.5 denotes the midway between the lowest and the highest possible scores. Error bars are SEM.

**Figure 2.**
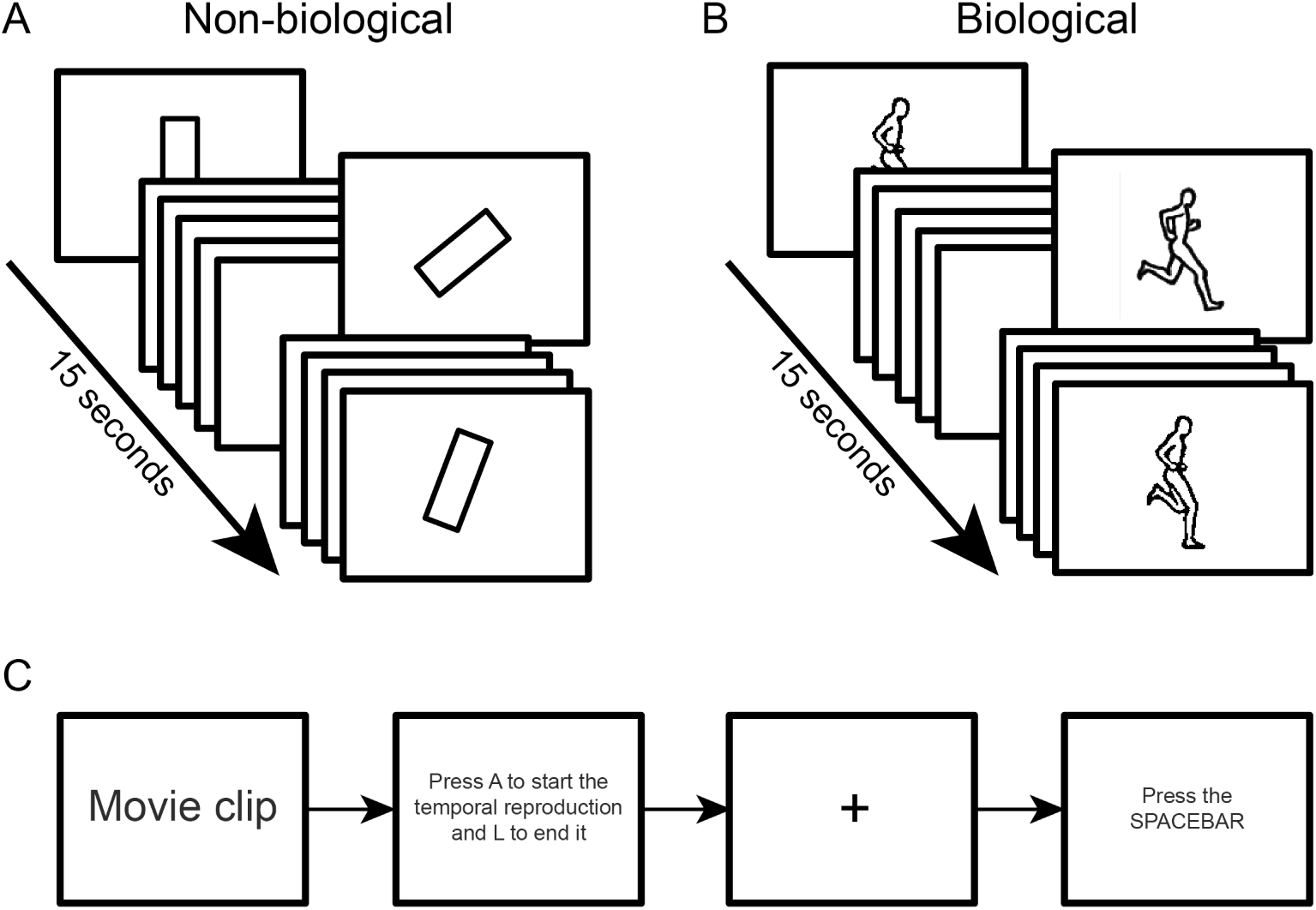
Task for Experiments II and III. Non-biological (A) and biological (B) movie clips were played for 15 seconds. C: Participants were instructed to press A to start the interval reproduction, and to press L to end it. Next, a press on the spacebar started the next trial.

Figure 1B shows average scores for how natural participants judged each stimulus type and frame rate to be. On average, biological and non-biological stimuli were judged as “natural” (close to zero) when presented at 6.4 or 12.8 fps, and as “non-natural” (close to −1) when presented at other frame rates. For the biological stimuli, mean (± SEM) estimates for plausibility were −1.00 (± 0.00), −1.00 (± 0.00), −0.5 (± 0.10), −0.14 (± 0.05), and −0.94 (± 0.04) for frame rates 0.2, 0.4, 6.4, 12.8, and 25.6, respectively. For the non-biological stimuli, mean (± SEM) estimates for plausibility were −1.00 (±0.00), −1.00 (± 0.00), −0.35 (± 0.11), −0.26 (± 0.09), and −0.94 (±0.05) for frame rates 0.2, 0.4, 6.4, 12.8, and 25.6, respectively. A GLM analysis with two factors, type of stimulus (biological and non-biological) and frame rate (0.2; 0.4; 6.4; 12.8, and 25.6 fps) showed an effect of frame rate (F(4,64) = 51.32, p < 0.001), no effect of type of stimulus (F(1,16) = 0.05, p = 0.835) and no type of stimulus x frame rate interaction (F(4,64) = 1.45, p = 0.226).

### Discussion

Experiment I sought to determine how much movement participants attributed to each movie clip, as well as how natural movement seemed to be at each frame rate. In accordance to previous studies [36], results showed that perceived speed positively correlated with frame rate, regardless of the type of stimulus (biological or non-biological, Figure 1A). Importantly, Experiment I determined that at intermediate frame rates (6.4 and 12.8 fps) movements seemed natural to participants.

Next, the goal was to assess the effects of frame rate and type of stimulus on temporal perception, and to describe any correlations to the assessments provided by Experiment I.

## Experiment II – Prospective temporal reproduction

In Experiment II, participants performed a prospective temporal reproduction task for the 15 seconds movie clips. Based on the speed-dependent temporal distortions previously reported, and given the results from Experiment I, we expected that movie clips played at higher frame rates would lead to an overestimation of their durations compared to movie clips played at lower frame rates, regardless of their biological or non-biological representation. We also expected the duration of movie clips of biological motion to be perceived differently from non-biological motion movie clips.

### Participants

Fifteen healthy undergraduate students participated in the second experiment (7 men, mean age: 23.2±3.4 years).

### Procedure

Participants performed the time reproduction task (Figure 2; see General materials and methods for the description of the task).

### Results

Figure 3 shows mean reproduced intervals for the two types of stimuli across the different frame rates. For the biological stimuli, mean (± SEM) reproduced intervals were 14.63 s (± 0.20), 14.82 s (± 0.34), 16.10 s (± 0.50), 16.04 s (± 0.50), and 15.91 s (± 0.46) for frame rates 0.2, 0.4, 6.4, 12.8, and 25.6, respectively. For the non-biological stimuli, mean (± SEM) reproduced intervals were 14.56 s (±0.36), 14.82 s (± 0.34), 16.10 s (± 0.50), 16.04 s (± 0.50), and 15.58 s (±0.48) for frame rates 0.2, 0.4, 6.4, 12.8, and 25.6, respectively.

**Figure 3.**
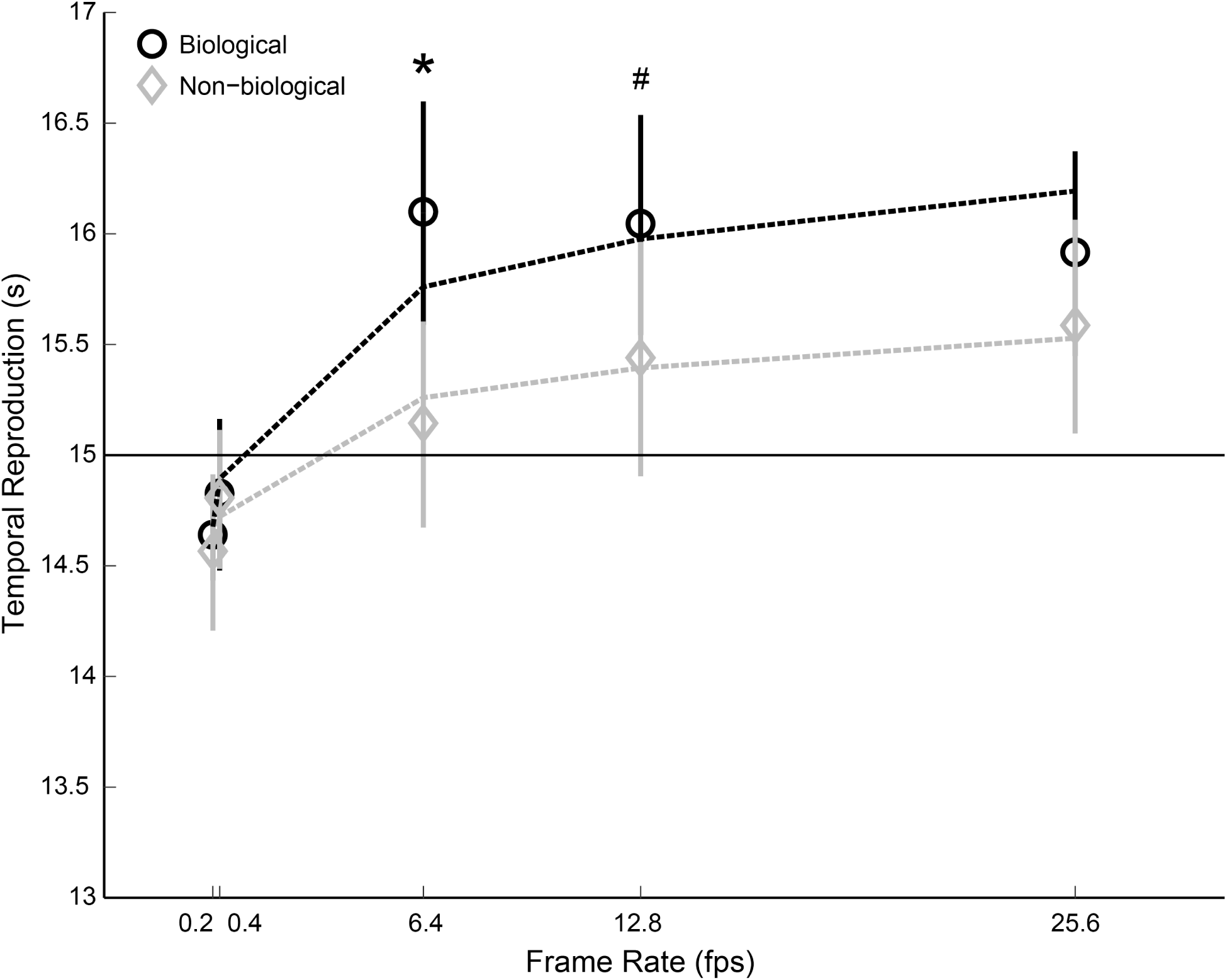
Experiment II, temporal reproductions. Mean reproduced intervals for biological (black) and non-biological (gray) movie clips as a function of frame rate. Dashed lines represent the log-linear fits to the data. Error bars are SEM.

A GLM analysis with two factors, type of stimulus (biological and non-biological) and frame rate (0.2; 0.4; 6.4; 12.8, and 25.6 fps) showed an effect of frame rate (F(4,56) = 5.63, p = 0.001), of type of stimulus (F(1,14) = 6.59, p = 0.022) and a significant type of stimulus × frame rate interaction (F(4,56) = 2.95, p = 0.028). A Bonferroni-corrected Wilcoxon test for paired samples indicated significant differences between biological and non-biological stimuli for 6.4 fps (adjusted α = 0.01; Z = 2.61, p = 0.009) and a marginally significant difference for 12.8 fps (adjusted α = 0.01; Z = 2.50, p = 0.012).

To further describe the influence of the type of stimulus and the different frame rates on temporal estimation, a log-linear model (Equation 1) was adjusted to individual data for the biological and non-biological conditions:

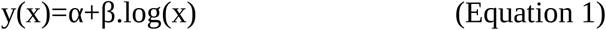

where α indicates initial rate and β indicates the slope. The best individual parameters, averaged across participants, were α = 14.9 and β = 0.19 (r^2^ = 0.96) for the non-biological condition, and α = 15.2 and β = 0.31 (r^2^ = 0.90) for the biological condition. A Wilcoxon test for paired samples showed no significant differences for the α parameter between conditions (Z = 1.60, p = 0.109), but a significant difference for the β parameter between the two conditions (Z = 2.33, p = 0.020).

### Discussion

Experiment II sought to assess the effects of frame rate and type of stimulus on temporal perception. Results showed that, as frame rate increased, so did the estimated durations for all movie clips, even though their real durations were kept constant at 15 seconds. This temporal illusion correlated with amount of movement perceived (Figure 1A), and suggests that amount of movement might mediate this illusion.

In addition, Experiment II confirmed differences in temporal perception for biological vs. non-biological stimuli, as previously reported, but only at frame rates for which movement was judged to be natural. This is an important addition to previous studies as it suggests that human representations of movement are necessary but not sufficient to elicit temporal representations that differ from non-human stimuli: human movement must also seem plausible.

Next, the goal was to reproduce the observed modulations of frame rate and type of stimulus on temporal perception while simultaneously assessing hemodynamic cortical activity, which has been previously shown to reflect biological vs. non-biological representations of movement.

## Experiment III – Prospective temporal reproduction and fNIRS

In Experiment III, participants performed the same prospective temporal reproduction task as in Experiment II. However, concomitant hemodynamic cortical responses over the bilateral STS were recorded using fNIRS. According to previous studies, we expected higher hemodynamic responses in the STS to smooth biological motion stimuli than to non-biological ones.

### Participants

The same participants from Experiment I also participated in the third experiment.

### Procedure

After filling out the questionnaires for Experiment I, participants stayed in the lab for Experiment III. They performed the same time reproduction task as in Experiment II, with only one difference: after pressing the L key to end the interval reproduction, a screen with a white cross on a black background was presented for 15 seconds. Once the 15 seconds had elapsed, the next trial began (i.e., there were no lever presses and, therefore, the time between trials was kept constant at 15 seconds).

### fNIRS acquisition

Participants performed the prospective temporal reproduction task while physiological recordings were obtained by functional Near Infrared Spectroscopy. Hemodynamic signal was acquired with a NIRScout 16 × 16 S/N: 073, NIRx hardware, and recorded by NIRx NIRStar 14-2 software. The regions of interest (ROIs) were set over the Superior Temporal Sulcus and Middle Temporal Gyrus (Figure 5A). Bilateral coordinates of each area were defined based on fMRI studies [32, 33, 34, 35, 37] and converted from Talairach coordinates to 10-10 NIRS channel system [38]. The total setup was formed by 43 channels (14 sources and 16 detectors) and the channels outside the regions of interest were used to perform a global signal regression.

**Figure 4.**
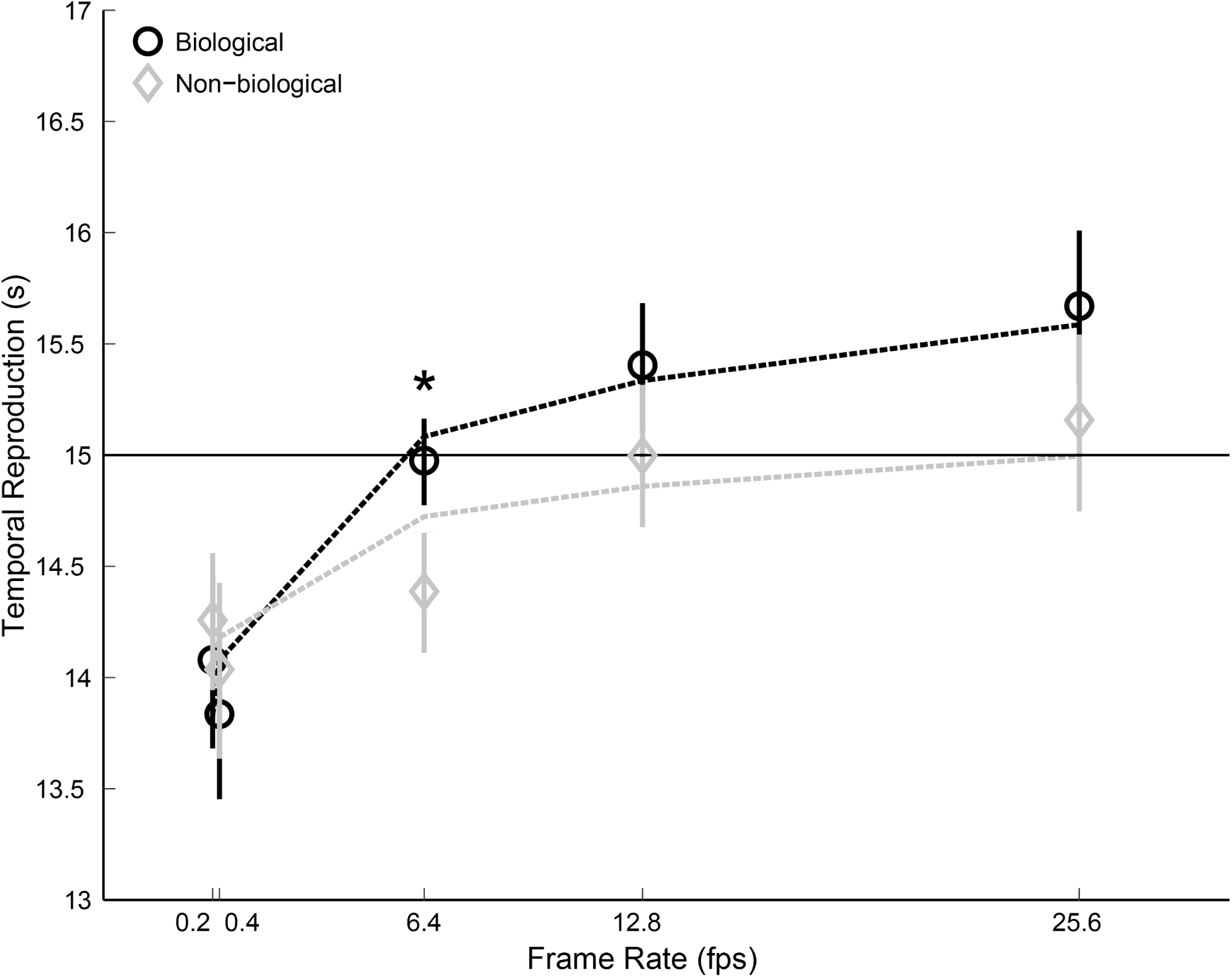
Experiment III, temporal reproductions. Mean reproduced intervals for biological (black) and non-biological (gray) movie clips as a function of frame rate. Dashed lines represent the log-linear fits to the data. Error bars are SEM.

**Figure 5.**
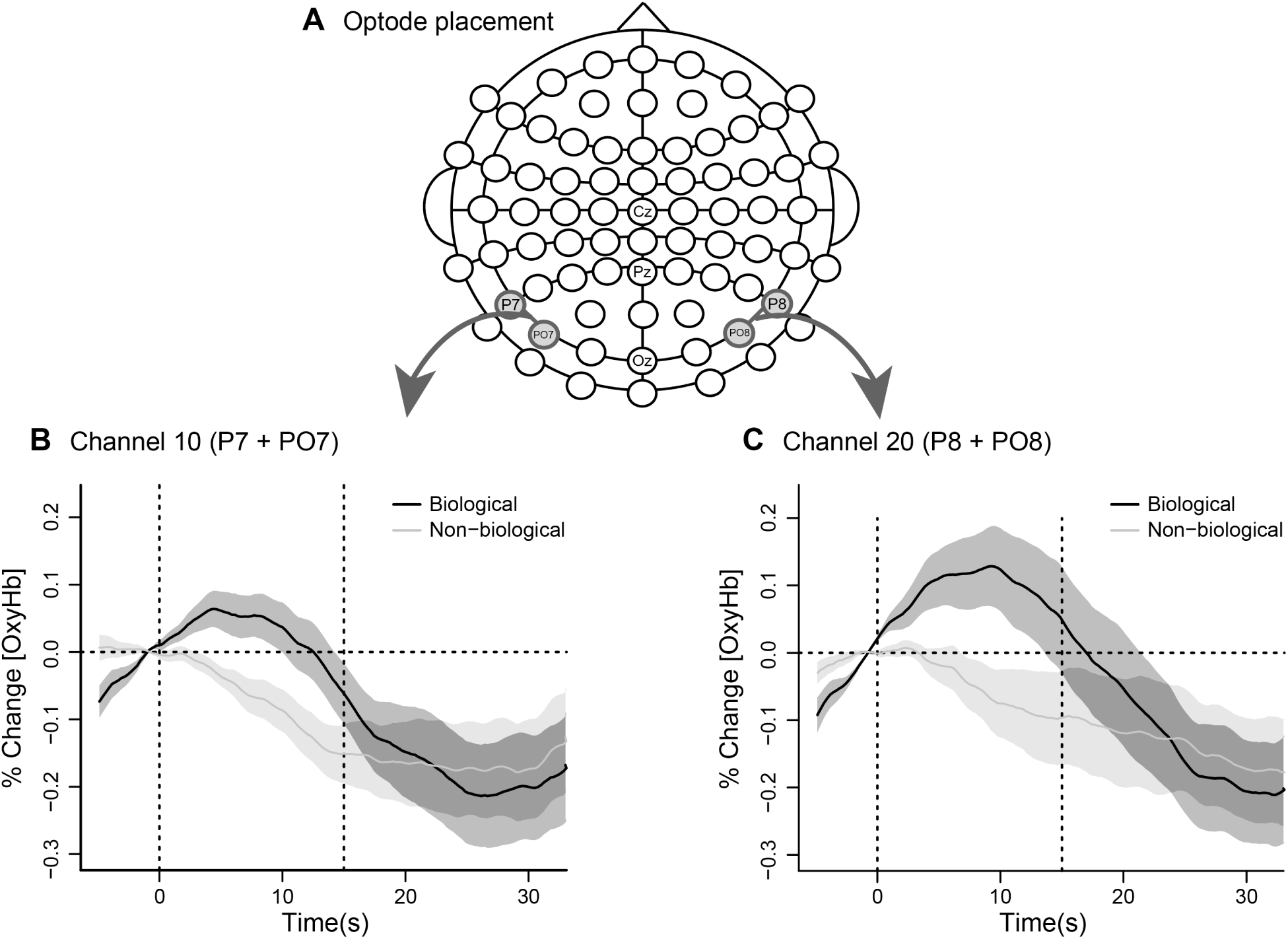
Experiment III, fNIRS cap configuration and channel time series. (A) Optodes placement for channel 10 (left side) and channel 20 (right side). (B) Time series of % change of O2Hb around the time during which the movie clips were played (0 – 15s, vertical dotted lines) for channel 10 and (C) for channel 20. Lines are means and shaded areas are SEM.

### fNIRS data analysis

Raw optical density data were imported and converted into relative concentration changes of oxy-Hb [O2Hb] and deoxy-Hb [HHb] according to the modified Beer-Lambert Law [39] in the nirsLAB analysis software. All data were imported to R for further analysis. A third degree Savitzky-Golay band-pass filter was applied [40]. Regarding quality control, channels was excluded based on the correlation between O2Hb and HHb as described by Cui, Bray and Reiss [41]. Hence, channels which presented a positive correlation between O2Hb and HHb during a complete block were excluded from further analyses. Participants with more than 70% of channels with a positive correlation between O2Hb and HHb were also excluded from further analyses. Following steps included a multi-channel (global) signal regression that used the median of all channels [42], detrending, normalization, and block averaging. Individual blocks were segmented according to the individual triggers. A block consisted of the signal recorded during the 15 s of stimulus presentation, a variable duration from time reproduction task, and a 15 s resting period.

Descriptive and inferential analyses focused only on the periods in which the behavioral data showed differences between biological and non-biological stimuli. Mean and standard error for the O2Hb signal during the 15-sec movie clips are reported for channels 10 and 20. They were first calculated individually (median signal), and then averaged across participants (mean of medians). A Mann-Whitney-Wilcoxon test was then used for statistical comparisons between plausible biological and non-biological medians.

### Behavioral Results

Figure 4 shows average reproduced intervals across participants for the two types of stimuli and the different frame rates. For the biological stimuli, mean (± SEM) reproduced intervals were 14.07 s (± 0.10), 13.83 s (± 0.09), 14.96 s (± 0.05), 15.39 s (± 0.07), and 15.66 s (± 0.09) for frame rates 0.2, 0.4, 6.4, 12.8, and 25.6, respectively. For the non-biological stimuli, mean (± SEM) reproduced intervals were 14.25 s (±0.07), 14.03 s (± 0.10), 14.38 s (± 0.06), 14.99 s (± 0.08), and 15.14 s (±0.10) for frame rates 0.2, 0.4, 6.4, 12.8, and 25.6, respectively.

A GLM analysis with two factors, type of stimulus (biological and non-biological) and frame rate (0.2; 0.4; 6.4; 12.8, and 25.6 fps) showed an effect of frame rate (F(4,64) = 9.59, p < 0.001), type of stimulus (F(1,16) = 6.96, p = 0.018) and a significant type of stimulus × frame rate interaction (F(4,64) = 3.30, p = 0.016). A Bonferroni-corrected t-test for paired samples indicated a marginal difference between biological and non-biological stimuli for 6.4 fps (adjusted α = 0.01; Z = 2.43, p = 0.012).

Similar to Experiment II, to further describe the influence of the type of stimulus and the different frame rates on temporal estimation, a log-linear model (Equation 1) was adjusted to individual data for the biological and non-biological conditions. The best individual parameters, averaged across participants, were α = 14.36 and β = 0.19 (r^2^ = 0.76) for the non-biological condition, and α = 14.41 and β = 0.36 (r^2^ = 0.94) for the biological condition. A Wilcoxon test for paired samples showed no significant differences for the α parameter between conditions (Z = 0.56, p = 0.560), but a significant difference for the β parameter between the two conditions (Z = 2.53, p = 0.010).

### fNIRS results

Figure 5B shows the time series variations for the O2Hb signal during the 15-s movie clips recorded for channel 10 (left hemisphere). For the biological stimuli, the mean signal (and SEM) was 0.041 (± 0.030), while for non-biological stimuli it was −0.063 (± 0.023). A paired two sample Mann-Whitney-Wilcoxon test revealed a significant difference between them (Z = 2.341, p = 0.019).

For channel 20 (right hemisphere, Figure 5C), mean O2Hb signal (and SEM) during the 15-s movie clips were 0.109 (± 0.052) and −0.046 (± 0.036) for biological and non-biological stimuli presentation, respectively. A paired two sample Mann-Whitney-Wilcoxon between plausible biological and non-biological means showed marginally significant differences between them (Z = 1.922, p = 0.055).

### Discussion

In Experiment III, while participants performed the prospective temporal reproduction task, hemodynamic cortical responses over the bilateral STS were recorded using fNIRS. As hypothesized, we observed higher hemodynamic responses in the STS to smooth biological motion stimuli than to non-biological ones. This finding is important for two reasons. First, it provides further evidence the STS might be involved in the processing of moving biological stimuli; and second, it reproduces the results observed from fMRI experiments with fNIRS, a more affordable and easy-to-access alternative technique for neurophysiological data acquisition.

## General Discussion

The present study assessed possible differences in temporal perception of movie clips composed of biological or non-biological stimuli as a function of frame rate. Previous studies have only indirectly investigated the effect of such manipulations on temporal reproduction [43]. Here, we kept the interval to-be-reproduced constant (15 seconds); explicitly manipulated the type of stimulus presented (biological or non-biological); and manipulated frame rates in a complete within-subjects factorial design.

Numerous studies have investigated behavioral differences between biological and non-biological motion stimuli. For example, dynamic point-light displays have been used to describe biological motion [44] and are able to infer gender [45] and even individual identity [46]. However, the recognition of a human body in such representations becomes more difficult if those point-lights move at low speeds or become static, demanding other visual clues such as the body outline, for its proper identification. To our knowledge no study has yet employed explicit representations of the human body, such as the human body outline, to assess its temporal properties in comparison to non-biological stimuli.

In accordance with previous studies [7-9], our results showed that biological and non-biological 15-s movie clips played at high frame rates (6.4 fps and higher) led to longer reproduced intervals as compared to movie clips played at low frame rates (0.2 and 0.4 fps). Increases in frame rate also led to increases in the amount of movement perceived, as judged by an independent group of participants.

Kanai and colleagues used a square moving on the computer screen at different speeds (degrees per second) and for different periods of time (from 200-1000 ms) [8]. Participants were required to reproduce the duration of movement of the square regardless of movement speed. A four-parameter model adjusted to the data indicated a plateau (i.e., ceiling effect) in the interval reproduced after movement reached a certain speed. In our study temporal reproduction data were fitted with a simplified two-parameter log-linear model. Although interval durations were considerably higher than those used by Kanai, adjusted parameters also showed a rapid increase in temporal reproduction for the set of lower frame rates (between 0.2, 0.4 and 6.4 fps) followed by a plateau after 6.4 fps for both biological and non-biological stimuli. These results can be interpreted as meaning that that pattern of temporal reproduction is not dependent on the range of intervals used.

More interestingly, movie clips of biological stimuli were overestimated compared to movie clips of non-biological stimuli when played at frame rates judged to be natural (6.4 and 12.8 fps), but not when played either too quickly (25.6 fps) or slowly (0.2 and 0.4 fps). These differences cannot be explained by possible differences between stimulus types in terms of perceived amount of movement or plausibility (i.e., how natural they appeared to be), as an independent group of participants explicitly judged those characteristics as similar.

A possible explanation, however, deals with the possibility that the neuroanatomical pathway for the processing of the visual information differs according to frame rate. For low frame rates (0.2 and 0.4 fps), processing may follow the ventral stream. This is because the dynamic information of movement is scarce in this condition, so the shapes of the figures may stand out, leading to a bottom-up processing of visual information. On the other hand, at intermediate frame rates (6.4 and 12.8 fps) in which the movement seems smooth and plausible, visual processing via both the ventral and dorsal streams is likely. In this case, top-down processing may overcome bottom-up processing specifically for the biological motion stimuli, as the movement of human silhouette becomes predictable. Finally, at the highest frame rate (25.6 fps), although the perceived smooth movement may require ventral and dorsal stream integration for visual processing, movement at that high rate was not rated as plausible. Therefore, top-down processing of such visual information may not be specific for the biological motion stimuli.

These results conflict with those reported by Gavazzi and colleagues, which suggested overestimation of time intervals by non-biological stimuli compared to biological stimuli [43]. In that study, however, intervals varied from 400-1900 ms, and the stimuli consisted of a dot that moved on a vertical axis on a computer screen, and travelled at either an increasing then decreasing acceleration (following the trajectory and kinematic of a person raising his/her arm; biological condition), or at a constant speed (non-biological condition). Since the study did not focus on differences in temporal perception between biological and non-biological motion as a function of speed of movement, interval duration and movement speed covaried in their study. That is, intervals were made shorter by increasing the speed of the moving dot. In our study, we varied frame rate while keeping the interval constant, and used a more naturalistic stimulus (human silhouette performing a familiar action) to represent biological motion.

The use of a human silhouette may have elicited some properties of embodiment, as previously reported [16]. Indeed, people are able to make inferences about perceived movement from biological stimuli based on their own motor skills, but not from non-biological motion stimuli [22-26]. It is possible that the difference herein reported in temporal reproduction at natural speeds between biological and non-biological stimuli may be supported by different corresponding underlying neural and cognitive processes. The perception of biological motion relies more heavily on complex and continuous bottom up and top down processes [25, 26] as compared to non-biological motion. Additionally, several neuroimaging studies have suggested that the processing of visual information related to biological and non-biological motion rely partially on different brain areas and networks. For example, the Superior Temporal Sulcus (STS) has been shown to increase activity for biological motion stimuli, while for non-biological stimuli the Middle Temporal Gyrus (MTG) seems to be more activated [29-33]. Alternatively, considering that former studies have reported distinct loci for visual processing of biological and non-biological motion stimuli, it is possible to hypothesize that those loci recruit processing from different networks, and so, certain time processing mechanisms involved in temporal perception of biological motion stimuli would be different from time processing mechanisms involved in temporal perception of non-biological motion stimuli.

Hemodynamics results have shown an increased activity in channels of interest during observation of biological motion stimuli compared to non-biological motion stimuli. This result is in accordance with our initial hypothesis, since the channels placement covered the area over STS. More interesting is the relationship between the differences in time perceived for biological and non-biological stimuli and the hemodynamic activity during the observation of biological and non-biological stimuli. Although it is still not possible to affirm that the hemodynamic activity in STS modulates the perceived time for biological motion stimuli, it may be informative since it could suggest different networks involved in visual and temporal perception of biological and non-biological stimuli.

In general, our study adds to the evidence that our perception of the duration of events is dependent on the nature of the event being timed and, more specifically, that naturalistic and plausible biological moving stimuli lead to a temporal overestimation compared to non-biological stimuli or implausible biological stimuli. Moreover, we have described the corresponding underlying hemodynamic activity of STS during those estimations using fNIRS, a less costly and more portable technique for neurophysiological data acquisition.

## Acknowledgments

This study was funded by Universidade Federal do ABC (UFABC). The authors would like to thank the members of the Timing and Cognition Laboratory at UFABC (http://neuro.ufabc.edu.br/timing/) for useful discussions and suggestions on this study. MSC is affiliated to Instituto Nacional de Ciência e Tecnologia sobre Comportamento, Cognição e Ensino, supported by the Brazilian National Research Council (CNPq, Grant # 465686/2014-1) and the São Paulo Research Foundation (Grant # 2014/50909-8).

